# Strong ‘functional’ divergence of tropical reef fish assemblages along the global diversity gradient

**DOI:** 10.1101/2020.09.18.303008

**Authors:** V Parravicini, MG Bender, S Villéger, F Leprieur, L Pellissier, GFA Donati, SR Floeter, EL Rezende, D Mouillot, M Kulbicki

## Abstract

Coral reefs are experiencing declines due to climate change and local human impacts. While at local scale biodiversity loss induces shifts in community structure, previous biogeographical analyses recorded consistent taxonomic structure of fish communities across global coral reefs. This suggest that regional communities represent a random subset of the global species and traits pool, whatever their species richness. Using distributional data on 3,586 fish species and latest advances in species distribution models we show that the global distribution of reef fishes is influenced by two major traits (body size and diet) and produces a strong divergence in the trait structure of assemblages across the biodiversity gradient. This divergence is best explained by the isolation of reefs during past unfavorable climatic conditions and highlights the risk of a global community re-organization if the ongoing climate-induced reef fragmentation is not halted.

## Background

Coral reefs are the most diverse marine ecosystems and show steep biodiversity gradients in numerous taxa that support a multitude of ecological functions and ecosystem services [1–5]. Concern has globally emerged about the future of this system as reefs are threatened by the combination of habitat loss and degradation due to climate change and local human impact such as overfishing [6–8]. Coral reef biodiversity is expected to be increasingly exposed to these effects. Coral bleaching may become an annual phenomenon for most coral reefs in less than 20 years [9] and fishing pressure is increasing with little capacity to move towards more sustainable practices [10]. In this context, identifying the mechanisms associated with community assembly and disassembly across the species richness gradient is important to elucidate potential effects of biodiversity erosion on this system.

On coral reefs, an apparent paradox calls to a closer examination of the role of the eco-evolutionary processes that determine community structure. While it is widely documented that biodiversity loss may produce severe shifts in the taxonomic and traits structure of communities at local scale [11– 14], biogeographical analyses suggested a remarkably consistent taxonomic structure across the globe for both corals and fish [15,16]. This would mean that despite strong evolutionary [17–19], environmental [2,4,20] and geographic drivers [5,21], the relative importance of species families at regional scale generally would not deviate much from a random subset of the global species pool. However, biogeographical patterns expressed solely in taxonomic terms may not capture the breath and distribution of species’ traits that are potentially correlated with ecosystem functioning [22]. Body size, for instance, is associated to a number of fish ecological characteristic, including colonization capacity, which may limit the distribution of species in isolated locations [23,24], especially when the surface of tropical reefs was strongly reduced during the coldest periods of the Quaternary [25]. At the same time, species diet may also determine the distribution of species according to resources availability and evolutionary mechanisms [26,27].

In this study we examined the prevalence of two important species traits for tropical reef fishes, namely body size and diet [27], across the global biodiversity gradient. Then, we used the latest modeling approaches to evaluate whether contemporary or historical factors best explained observed patterns. We hypothesize that contrary to the previously documented taxonomic pattern [15], trait structure of reef fish significantly vary across the biodiversity gradient. Specifically, the prevalence of body size classes in regional assemblages of tropical reef fishes would be primarily explained by the isolation from Quaternary refugia, as already showed for species richness [25] with a greater proportion of large-bodied species in the most isolated reef areas. Then, the prevalence of certain diet categories would be more strongly influenced by the area of tropical reefs, which can reflect the level of resources availability [28].

## Materials and Methods

### Distributional and trait data on reef fishes

We used a global database on tropical coastal fish occurrences previously compiled by Parravicini et al. [29]. These data consist in the global distribution of 6,316 fish species defined according to a grid of 5°×5° resolution, corresponding to approximately 550×550 km at the equator. Out of these species, we used distributional data for the 3,586 species considered in the fish phylogeny.

We focused our analysis upon two major species ecological traits: maximum body size and diet. Maximum body size in cm was extracted from FishBase while species diet was defined according to eight trophic guilds defined according to the quantitative analysis of fish gut content data and reef fish phylogeny (see Parravicini et al. [30]). This analysis uses detailed gut content information for 615 reef fish species and their phylogenetic position to assign species to the following trophic guilds: herbivores, microvores and detritivores (HMD), planktivores, sessile invertivores, corallivores, microinvertivores, macroinvertivores, crustacivores, piscivores.

### Statistical analysis

In order to test whether the trait structure of reef fish assemblages was stable along the global biodiversity gradient we performed two multinomial Bayesian models that predict the probability for a species to belong to a certain trait category (either a body size class or a trophic guild) according to the local species richness. For this analysis, similarly to Mouillot et al.[16], we defined body size according to 6 categories: 0–7cm, 7.1–15cm, 15.1–30cm, 30.1–50cm, 50.1–80cm.

Bayesian models were performed using the *brms* package in R using a multinomial logit link function. The probability for a species to belong to a certain category (either size category or trophic guild) is computed as follows:

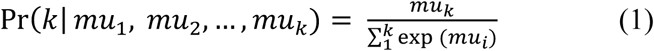

with *mu*_*k*_ defined as:

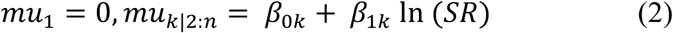

where *β*_*0k*_ is the intercept of the category-specific fixed effect, *β*_*1k*_ is the slope for the natural log transformed species richness (SR) for each category *k* (either body size classes or trophic guilds) and *n* is the number of categories (6 in the case of body size classes and 8 in the case of trophic guilds). For both body size and trophic guilds models we used uninformative priors and ran the model for four chains using 5,000 iterations per chain after 1,000 warmup iterations.

We then tested whether species body size and diet play a role in determining the spatial distribution of species. In other words, we tested whether the presence-absence of a species in a given location is due to the environmental characteristic of the location, the ecological characteristic of the species or the interaction between environment and the species traits. Understanding whether the environmental response of species is due to their ecological characteristics is known as the fourth-corner problem, that is the study of the environment–trait association using matrices of presence/absence across species, environmental data across locations and trait data across species. The matrix of environment–trait interaction coefficients is the fourth corner [31]. We used the generalized joint attribute modelling (GJAM) approach proposed by Clark et al. [32], which solves the fourth-corner problem accounting for the joint distribution of multiple species, thereby accounting for their potential interaction. In particular the GJAM approach was used to model the presence absence of the 3,586 species in each grid cell according to sea surface temperature, present coral reef area, present geographical isolation from coral reefs, coral reef area during the Quaternary and geographical isolation from coral reefs during the Quaternary. These variables were selected because they have already be shown as important correlated of species richness[4,25]. Mean SST (°C) for each grid cell was obtained from the Bio-ORACLE database at a resolution of 5 arcmin [33]. Estimates of coral reef area (km^2^) were obtained from data based on the Coral Reef Millennium Census project and available at: http://data.unep-wcmc.org/. Isolation estimates were calculated using a nearest neighbour approach. Isolation from present reefs, from reefs during the Quaternary and coral reef area during the Quaternary were obtained by Pellissier et al. [25]. For this analysis trophic guilds were treated as categorical variables, thus producing one slope for each guild-environmental variable combination, while maximum body size was treated as a continuous variable. Positive slope coefficients for the interaction between environmental variables and trophic guilds indicate that a certain environmental variable favours the presence of species that belong to a particular guild, while negative values indicate that a certain environmental variable favour the absence of species that belong to a particular guild. Instead, for maximum body size, positive slope coefficients for the interaction between body size and environmental variables indicate that that particular environmental variable favours the presence of large species and negative values indicate that the environmental variable favour the presence of smaller species.

## Results and Discussion

Contrary to the already known overall stability in the relative (taxonomic) contribution of fish families to local species richness, dominance of ecological traits markedly differs among reef fish assemblages at the global scale (Fig. 1). For instance, piscivores represent up to 20% of the species richness in the Tropical Eastern Pacific, while they do not exceed 12% in the Indo-Pacific (in the richest areas of the Indo-Australian Archipelago). Similarly, large species (>80cm in body size) constitute up to 40% of the total species richness in the Eastern Atlantic but only 6% in the Indo-Pacific.

**Figure 1.**
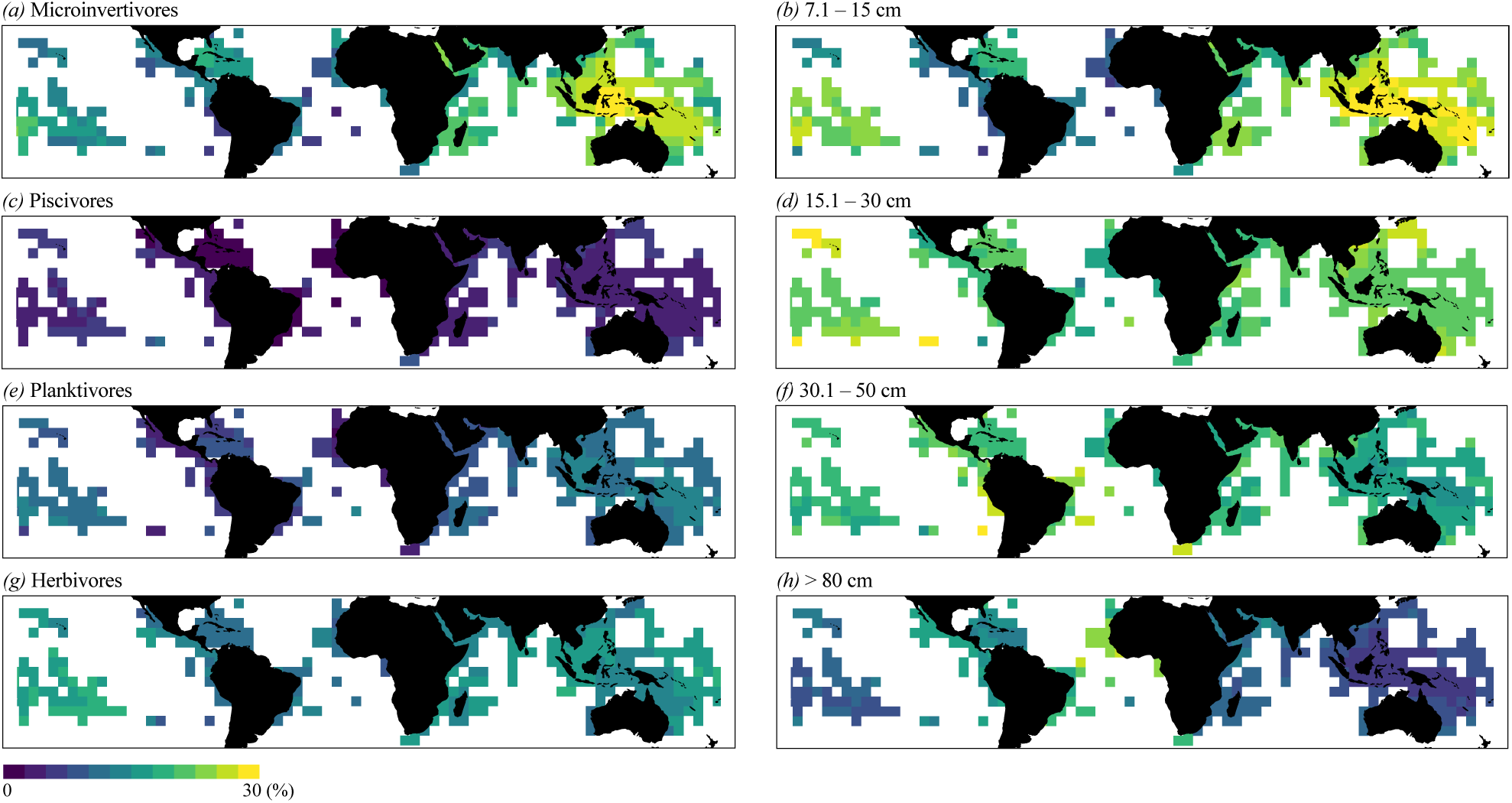
Global gradients in the proportion of species belonging to different trophic guilds and body size classes.

Small species tend to dominate species-rich areas in both the Indo-Pacific and the Atlantic, while larger species are proportionally dominant in more depauperate reefs. A similar pattern can be identified for diet categories. Micro-invertivores, planktivores and herbivores tend to dominate in the richest areas while poorer reefs appear to be dominated by piscivores, macro-invertivores and crustacivores.

In order to test whether regional assemblages can be considered a random subset of the global species pool we built two Bayesian multinomial models that quantify the probability of a species to belong to the different trait categories for both diet and maximum body size. Species body size was defined according to 6 categories as in Mouillot et al. [16] while diet was described according to the 8 categories recently proposed by Parravicini et al. [30] on the basis of assembled gut content data.

Our results highlight a remarkable divergence in the relative importance of traits across the species richness gradient for both maximum body size and species diet (Fig. 2). With the exception of few trait categories (i.e. sessile invertivores, corallivores and species of intermediate body size from 15 to 30cm) all the other trait categories displayed an uneven probability of occurrence across the global gradient in species richness. In poorer assemblages a species has higher probability of being large-bodied (>30cm) and either crustacivore or piscivore. On the other hand, in highly diverse locations, a species has the highest probability of being small-bodied (<15 cm) and having a microinvertivore diet.

**Figure 2.**
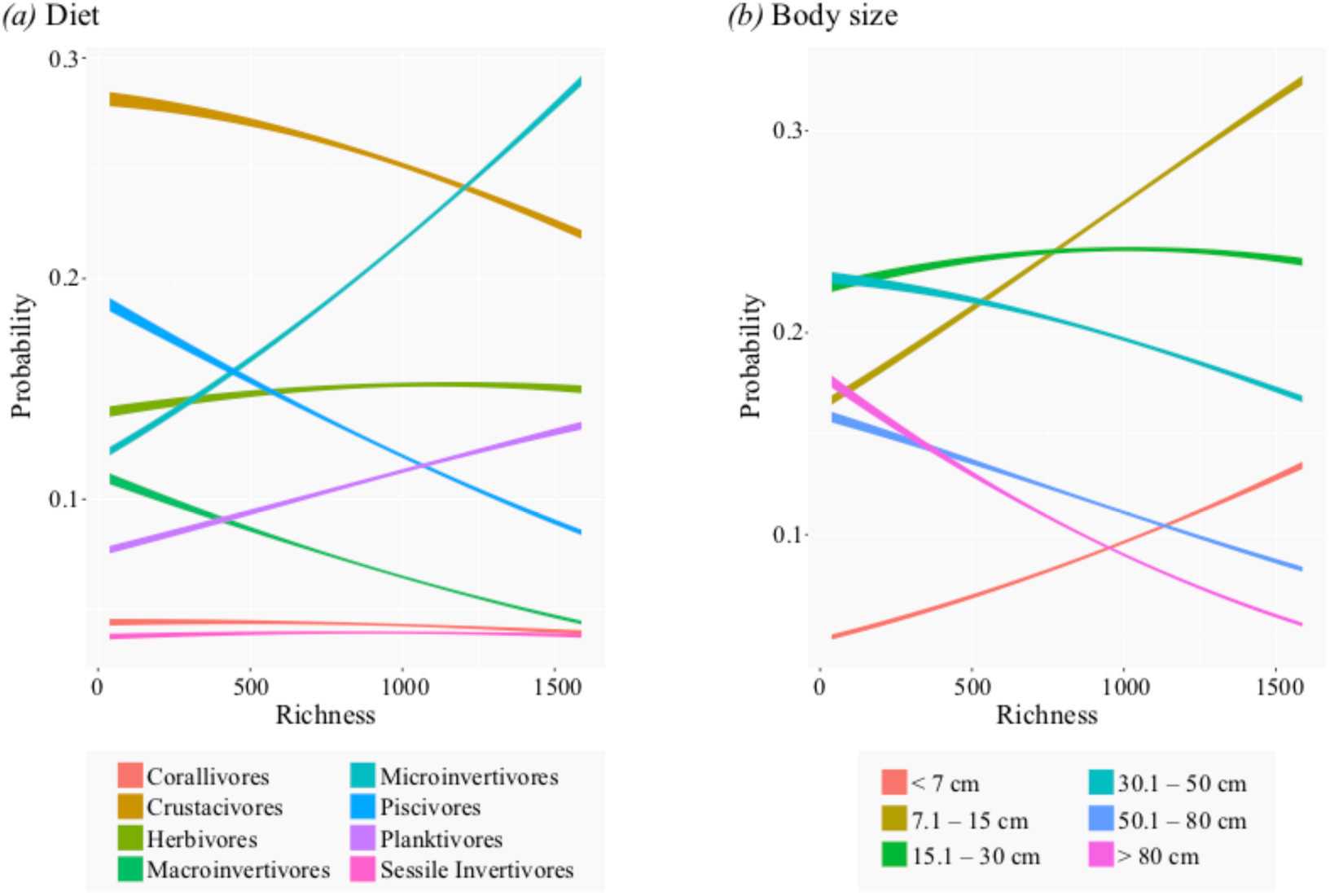
Marginal effects of species richness on the probability for a species to belong to different (a) trophic guilds or (b) body size classes.

Observing such a strong shift in the trait structure of assemblages across global coral reefs, we hypothesized that the same drivers of species richness may have interacted with certain traits across evolutionary time scale, thereby producing assemblages with markedly different trait structures. In order to test for such an interaction, we took advantage of the latest advances in species distribution modelling. Specifically, we used the same correlates of reef fish species richness (sea surface temperature, present reef area, past reef area, present geographical isolation and past geographical isolation) as in Pellissier et al. [25] and we built joint species distribution models using the GJAM modeling approach developed by Clark et al. [32]. We used such a technique to model the joint distribution of all the species according to present and past environmental variables, their ecological traits and the interaction between environmental variables and ecological traits. For this particular analysis body size was considered as a continuous variable; while species diet was described with the above mentioned eight categories.

The technique allowed us to identify a major role of ecological traits in determining species distribution (Fig. 3). In particular, both body size and diet strongly interacted with the isolation of coral reefs from refugia during the Quaternary. Indeed, our results suggest that past isolation caused by the Quartenary glaciations may have particularly favored large species, and among them, piscivores, crustacivores and macroinvertivores.

**Figure 3.**
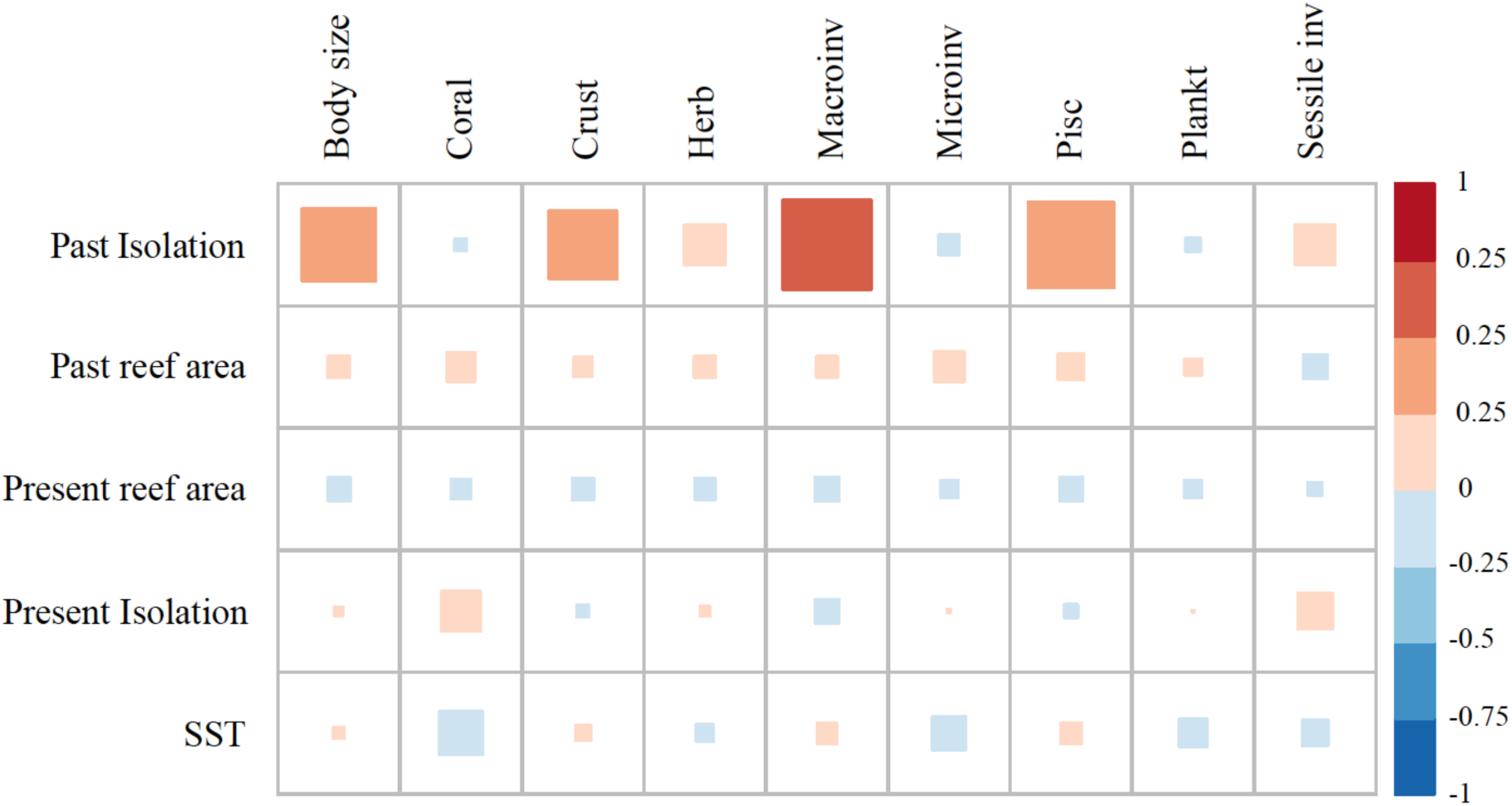
Slope coefficients returned by the generalized joint attribute model and referred to the interaction between species traits and environmental variables. Positive values correspond to a positive effect of the interaction in determining the presence of species while negative values indicate a negative effect on species presence. Body size is a continuous variable while diet is described according to 8 trophic guilds (Coral: corallivores, Crust: crustacivores, Herb: herbivores, Macroinv: macroivertebrate feeders, Microinv: microinvertebrate feeders, Pisc: piscivores, Plankt: planktivores, Sessile inv: sessile invertebrate feeders)

Overall, our results suggest that past environmental changes left a strong imprint on the trait structure of regional assemblages of reef fishes, and more particularly the isolation of reefs during unfavorable climatic events. During the Quaternary period, which was accounted for in our model, repeated periods of global cooling and warming caused coral reef to contract and expand [28]. Older isolation processes, not explicitly considered in our study, may also explain the observed prevalence of ecological traits globally. The historical biogeography of reef fishes [34] suggests that from the Oligocene onward, the Indo-Pacific experienced a history of connectivity, while the Eastern Pacific and the Atlantic were subjected to a history of isolation. The connectivity of the Indo-Pacific culminates during the Pliocene. In this favorable period coral reefs were expanding [17], and species originated in the IAA with subsequent movement toward the Central Pacific and the Indian Ocean. By contrast, during this period, the Tropical Eastern Pacific and the Atlantic Oceans were experiencing a contraction of their species pool probably due to extinction and increased isolation of their coral reefs[18]. In this context, our results support the hypothesis that large carnivores, with high colonization capacities and post-dispersal persistence abilities [23,35], had greater chances of survival or recolonization in the Atlantic and Tropical Eastern Pacific than smaller species. This process holds within oceanic basins with large carnivores characterizing depauperate locations that experienced extreme isolation in the past, or that are isolated from rich reef areas by barriers [34] or geographic distance [2,4,5]. Large species, showing high colonization capacity [23,36], may therefore have been favored in species-poor locations, presumably because of their ability to persist when facing unfavorable conditions or to better colonize after regional extinctions. By contrast, small species appear to have persisted better in locations with a high coral reef area, such as the IAA and the Caribbean. These biodiversity hotspots with large and less fragmented coral reefs have possibly favored the diversification (or at least the persistence, i.e. less extinctions) of small-bodied fish species that have comparatively higher population turnover rates, speciation rates [37] and therefore greater contribution to richness in these assemblages.

The spatial patterns reported here, although based on coarse categorization of species, still highlight two aspects of major concern when considering the present threats to coral reefs. First, we detected a variable trait structure of fish assemblages across the global biodiversity gradient, which suggests the potential for a functional re-organization in the case of regional species loss. Second, our findings suggest that the trait structure of reef fish assemblages at the regional scale reflects the interaction between species’ colonization capacity with reef isolation and fragmentation across unfavorable climatic periods. Overall, our analysis implies that ongoing coral reef fragmentation due to recent climate-induced global scale mortality of corals may alter key variables that contributed to the unique trait structure of global biodiversity hotspots.

Conservation actions cannot address the vulnerability of assemblages inherited from the past. Nevertheless, tailored and realm-specific conservation actions may be necessary to ensure connectivity under present conditions and future scenarios. Conservation efforts aimed to sustain coral reef connectivity will be essential to maintain populations and species, as well as ecosystem processes, by preventing functional alteration, triggered by the loss of species with limited colonization ability.

